# On the dimensionality of olfactory space

**DOI:** 10.1101/022103

**Authors:** Marcelo O. Magnasco, Andreas Keller, Leslie B. Vosshall

**Affiliations:** Laboratory of Mathematical Physics, The Rockefeller University, 1230 York Avenue, Box 212, New York, NY 10065 USA; Laboratory of Neurogenetics and Behavior, The Rockefeller University, 1230 York Avenue, Box 63, New York, NY 10065 USA; Howard Hughes Medical Institute, 1230 York Avenue, Box 63, New York, NY 10065 USA

## Abstract

We recently presented an estimate of the number of mutually discriminable olfactory stimuli at one trillion (*1*). Subjects were asked to sniff mixtures of molecules with increasing component overlap selected from a panel of 128 isointense structurally and perceptually diverse monomolecular odorants (*2*). We considered stimulus pairs discriminable when the majority of subjects could significantly discriminate them at p=0.05, a conventional statistical threshold given our sample size. From these empirical data, we estimated that human discriminative capacity exceeds one trillion olfactory stimuli. Several readers have pointed out that such extrapolations are sensitive to underlying assumptions about the chosen significance threshold (*3*) and the dimensionality of olfaction (*4*). It is important to note that any exponential function will be sensitive in this way, and the goal of our model was not to identify the exact number of discriminable olfactory stimuli, or even the exact mathematical bounds, but an estimate of the order of magnitude of human discriminatory power across a population of human subjects. This was not clearly stated in our paper, and we agree that contradictory references to a “lower limit” and an “upper bound” were confusing. The central argument in (*4*) is that our estimation method assumes that the dimensionality of olfactory space is large. We agree that the high-dimensional nature of olfaction is indeed an assumption, and we should have stated this explicitly in our paper (*1*). Even if we follow this logic of the models presented in (*4*), purely geometrical calculations show that our results hold if the dimensionality of olfactory representations is D≥25. The dimensionality of olfaction is a question of interest to everyone, and while we do not know for sure, all available evidence suggests that olfaction is a high-dimensional sense. The olfactory system is wired to keep information from the ~400 odorant receptors strictly separated, so it is plausible that olfaction operates at least in 400-dimensional space. This is an important topic of discussion in olfaction, and we welcome continued debate of the dimensionality of smell and how this impacts human olfactory perception.

## Introduction

One of the key open problems in olfaction research is to elucidate the global structure of olfactory perceptual space, including such basic features as its dimensionality and overall topology. Our ignorance about this space in olfaction compared to visual perceptual space is one of the main limitations in understanding the cognitive mechanisms of this sense. Compared to the situation in color vision, in which the one-dimensional stimulus space of the visual spectrum (390-700 nm), the spectral tuning of the receptors, and the three dimensional perceptual space (hue, saturation, brightness) are known, essentially none of the corresponding parameters is known in olfaction.

To begin to address some of these questions, we carried empirical experiments to probe the ability of humans to discriminate odor stimuli (*1*). This work required us to pick odor stimuli appropriate for this task, and we selected mixtures of molecules selected from a panel of 128 isointense monomolecular odorants that are structurally and perceptually diverse. This unbiased method of creating stimuli for olfactory psychophysics was developed by Noam Sobel’s group (*2*). An important advantage of these stimulus mixtures is that they do not create recognizable odor percepts that are readily characterized with semantic descriptors such as “rose,” or “lemon”, or “tobacco.” These unfamiliar scents allowed subjects to focus on a task of pure discrimination, without the distraction of putting words to familiar odors. We tested 13 types of mixtures, which differed in the number of molecules (10, 20, or 30), and the number of overlapping components. Subjects were presented with 3 vials, one of which was different from the other two, and were instructed to pick the odd stimulus. The difficulty of the discrimination task increased with the degree of component overlap between mixtures. From these data, we estimated that human discriminative capacity is large, exceeding one trillion olfactory stimuli. Working from our raw data, Gerkin and Castro and Meister (*4*), present alternate mathematical approaches leading to different extrapolations, which we discuss below.

Gerkin and Castro (*3*) write that “‘Significantly discriminable’ is a moving target.” They point out that the number of discriminable stimuli drops several orders of magnitude when the significance threshold is set to p=0.0005, and that the estimate would increase if more subjects and more stimuli were tested. This is true, and would hold for any similar dataset. For instance, a genome-wide association study designed to identify DNA variations that are linked to a disease could yield significant association of most of the genome or no gene at all, merely by adjusting the significance threshold down or up, respectively. The power to find such associations would rise at some rate merely by increasing the number of subjects genotyped, at the extreme every human on earth. These effects of significance threshold selection and sample size do not invalidate genome-wide association or our own work. What is required is a prudent and defensible statistical cutoff and sample size.

So on what basis is it appropriate to say that a pair of stimuli is discriminable? Is it enough when one subject can discriminate an odor pair? Or should we require that all subjects discriminate a given stimulus pair before we consider it discriminable? These are important questions to pose given the striking variability in the human sense of smell. The remarkable interindividual variability in human olfactory performance makes choosing either extreme problematic: even in our population of 26 subjects, performance ranged from just above chance to 71% correct. Arbitrarily setting the threshold of discrimination to stimuli that all subjects must be able to discriminate or that any one subject on earth can distinguish would indeed lead to arbitrarily small or large numbers, respectively. In the extrapolations carried out in our work, our goal was to derive an estimate between these two extremes, and we therefore considered stimuli discriminable when ≥50% of subjects correctly identified a mixture type. One can certainly quibble with our statistical assumptions, or the threshold we chose, but the mere fact that changing them will lead to different outcomes does not invalidate our conclusions, any more than it would invalidate the entire field of genome association studies.

Meister presents three models based on our empirical psychophysical data, and points out that our extrapolation fails in regimes of low dimensionality (*4*). This is true, and the implicit assumption that olfactory perceptual space is high-dimensional should have been explicitly stated in our paper. The central feature of the three models proposed by Meister is that at large enough distances, percepts begin to repeat. A small change would lead to many new percepts, while a larger change would begin to repeat those already smelled. The simplest nontrivial way in which the exponentially-growing space of combinations can fail to give exponentially many percepts is if the latter is a low-dimensional Euclidean space. This conclusion is grounded in the assumption that olfaction follows the low-dimensional rules of color perception. Extensive chemical, genetic, neurobiological, and psychophysical data indicate that this is unlikely to be true. Although the number of molecules with the chemical properties of odors stands in the billions (*5, 6*), there is at present no means to predict the olfactory percept of a molecule from its chemical structure. Further, olfaction is a synthetic sense, and single molecules can be blended into mixtures to yield novel percepts that are not a simple transitive property between the components. As a result we know essentially nothing about the boundaries or dimensionality of olfactory stimulus space. The odorant receptors that detect these molecules comprise the single largest family of genes in any vertebrate, with more than ~400 functional odorant receptors in humans (*7, 8*) and >1000 in rodents (*9, 10*). These receptors are not narrowly tuned like visual receptors. If each recognized a single cognate odorant, humans would be able to detect only 400 smells. Instead, each odorant receptor responds to multiple olfactory stimuli, and a given molecule activates multiple odorant receptors (*11-14*). This combinatorial coding logic leads to a dramatic increase in dimensionality. Finally, every effort is made to maintain information from the 400 receptors as separate channels (*15*) until they reach higher olfactory cortex, where the coding complexity is vastly increased by random selection of subsets of inputs (*16*).

The “dimensionality of olfaction” on which our result rests is the dimensionality of the representation, namely, the number of independent information channels conveying actionable information to the higher brain areas. What is this numbers for olfaction? The truth is we do not know either with certainty. But as we shall argue in the following sections, the primary sensory representation is about 400-dimensional for humans. In this paper, we first review some important distinctions between olfactory discrimination and the entirely different task of olfactory categorization. With these distinctions in hand, we review the extant biological knowledge on olfactory pathways germane to dimensionality of representations. Then we return to our work (*1*) to explain why the dimensionality of olfactory representations affects our estimate, and show that a Euclidean dimension of about 25 suffices to embed every stimulus in our combination space. Then we discuss which features of the models presented in (*4*) generate the underlying differences with our data, and we extend the RGB model into D-dimensional space to show that 20 dimensions suffice to generate the trillion percepts of our estimate without repetition.

## Odor categorization vs. odor discrimination

The biological question asked in our work (*1*) was whether certain olfactory stimuli can be distinguished from each other. The task was not to recognize or categorize or express an emotion about individual smells, nor to describe the nature of their difference, but solely to detect the existence of a difference, even if lacking any semantic label for it. The percept one gets when presented with a single stimulus is entirely different from the perception of a difference, however subtle, between consecutive presentations of two stimuli. Although the underlying logic of olfactory perception has been debated for centuries, our study was the first attempt to address the discriminative capacity of humans experimentally.

The experience we have when we smell an odor is complex (*17*). We can say how strong it is (intensity), how pleasant or unpleasant it is (valence), and what it smells like (categorization). A completely different task is to ask if two odors smell different (discrimination). The olfactory psychophysical tasks of categorization and discrimination are frequently confused. Because different proposals about the structure of the perceptual olfactory space may be based merely on differences in terminology, we define what these tasks attempt to measure and why they are completely different.

Prior to our work, it was generally “known” that humans can discriminate 10,000 odors. In 1927, Crocker and Henderson built a model of olfaction based on the assumption that odor space is four-dimensional, with the dimensions being “fragrant, or sweet”; “acid, or sour”; “burnt, or empyreumatic”; and “caprylic, or oenanthic” (goat-like)(*18*). These authors proposed that nine levels, from 0 to 8, along each dimension can be discriminated (*18*). This odor categorization system results in 9^4^ or 6561 discriminable odors, each identified by a four digit code. Rose scent, for example, has the code 6423 (fragrant=6, acidic=4, burnt=2, goat-like=3), and freshly roasted coffee the code 7683. In subsequent years, this number was rounded up to 10,000 and its purely theoretical basis was forgotten (*19*).

In the 1980s, Dravnieks expanded the essential olfactory qualities by using 146 descriptors rated along a 5 point scale (*20, 21*). These descriptors were developed for use by the flavor and fragrance industry, and represent only a fraction of all descriptors that could be used to profile odor quality. The same scent of a rose profiled as 6423 by Crocker and Henderson was expanded to the 5 Dravnieks descriptors that pertain to flowers (“Rose” “geranium leaves” “violets” “lavender” “floral”), each of which can be rated along a 5-point scale. But if one feels that different rose scents deserve their own category, for example that of a budding and of a blooming rose, or those of the thousands of different rose cultivars, then one can make the categorization more fine-grained. One can increase the resolution of this categorization by increasing the number of dimensions, by adding “fresh,” “flowery,” or “rose-like,” etc. as a category, or by increasing the number of levels into which each dimension is subdivided. However, no matter how fine-grained a categorization task is made, one cannot draw conclusions about discriminability from odor categories because they necessarily collapse perceptual space into the dimensions defined by semantic descriptors.

Another obvious problem is that odor categories are strongly influenced by language and culture. “Caprylic” (from the Latin caper = goat) was introduced as an important semantic label for an odor category by Aristotle, who presumably had frequent encounters with goat odors. It is less popular as a dimension of odor descriptor space in the 20^th^ century. The descriptors “vanilla” and “chocolate” were introduced into language after products with these odors were introduced into Europe. As they were more frequently encountered, and grew in popularity as smells (*22*), vanilla and chocolate reached an importance that justified a verbal label.

Odor categorization gives interesting insights into the connection between language and smell perception, and it is extremely important for all practical applications that are built on an orderly system of olfactory stimuli. In our paper (*1*), we abandoned the inherent constraints of using language to describe smell perception and instead probed perception directly. The structure of olfactory space has traditionally been investigated using single odorous molecules, yet smells encountered in the environment are almost never single molecules. The perceived odor of a rose is the consequence of mixtures of hundreds of different molecules activating our olfactory sensory neurons. Olfactory spaces based on monomolecular odors therefore cover only a very small percentage of all the possible olfactory percepts. To overcome this limitation, we used mixtures of 10 to 30 different odorous molecules diluted to roughly equal intensity. These isointense mixtures were only recently introduced to olfactory psychophysics by Noam Sobel and his group, but they have already enabled impressive progress in quantitative olfactory psychophysics (*23*). Our work builds on this progress.

## Dimensions of a sensory representation

The number of statistically independent numbers gathered by the sensory receptors from the environment can be thought of as the dimensionality of a sense. This number is then affected by subsequent neural transformations into higher order brain areas. This dimensionality should not be confused with the fact that the receptors may be organized according to some set of parameters. These two numbers are quite distinct.

One example is the one-megapixel grayscale camera, in which the pixels are optically independent, which produces data whose dimensionality is one million. However, its pixels are arranged on a two-dimensional surface, so we say that the sensor array is organized in two dimensions with “2D topology” because its million sensors are parameterized by a two-dimensional index (x,y). One million is a very different number from two, not just in magnitude, but because the former can be easily changed by upgrading to a three-megapixel camera, while the latter is dictated by the physics of the sensing element.

Color cones convey 3-dimensional color information, but the cones themselves are labeled by a single parameter, wavelength, which is one-dimensional. Even in pentachromatic birds such as pigeons, their parameter is still one-dimensional even though their hyperspectral color space is five-dimensional, because their opsins are still organized by wavelength. Combining color with space, the dimensionality of the representation is the product of the dimensions, while the topology is the sum of the dimensions. A color photograph 4000 pixels in width by 3000 pixels in height is an array of 4000 × 3000 × 3 numbers, and is therefore a vector in a 36-million-dimensional space (12 megapixels times three colors) organized in three dimensions (two space plus one color).

Subsequent transformations of the neural representation as the information flows to other brain areas can reduce the dimensionality of the representation by collapsing information from many channels into one. This can and does happen for many reasons, and bitter taste is an instructive example. Each mammalian bitter cell expresses all of the ~30 bitter taste receptors (*24*) and is wired into a gustotopic map of bitterness in the brain (*25, 26*). This allows the cells to sense broadly a large stimulus space of bitter molecules, but collapses the ~30-dimensional space of bitters being sensed into a single dimension, the activation of bitter cells. This is an irreversible transformation.

## Chemical and biological for the high-dimensionality of olfaction

What is the current estimate for the number of olfactory stimuli that humans can detect and how are these arranged in stimulus space? For a molecule to have a perceptible odor, it must have a vapor pressure sufficiently low to be volatile and inhaled into the nose (*27*). Generally molecules with a molecular weight <500 Da meet this criterion, and of the 10^60^ such compounds (*6*) there are over 166 billion possible molecules of 17 or fewer atoms of C, N, O, S, and halogens (*5*). Among these molecules, even those with very similar chemical structure can be discriminated by humans (*28-30*). Conversely, small changes in chemical structure have dramatic effects on the ability of the compound to activate a given receptor (*12*). Recently, efforts have been made to relate chemical structure to perception (*23, 31*), and this work relies on over 4,800 chemical descriptors that can be used to profile an olfactory stimulus (*32*). Despite more than a century of attempts to relate chemical structure to percept, we still have no reliable means to predict what a molecule will smell like, although recent work from Noam Sobel and colleagues has begun to provide insights (*23, 33*). More profoundly, there is no information about how all possible olfactory stimuli are arranged in stimulus space, and what the boundaries of our perception are. There is not yet a “visible spectrum” for olfaction that defines the range of detectible smells.

Turning to the biological evidence for olfactory dimensions, we note that the stimulus tuning and wiring of sensory input defines the organizational logic of the resulting percept. All available evidence points to this sense being high-dimensional. There are about 400 functional olfactory receptors in the human genome (*7, 8*) and 1000 in the rodent genome (*10, 34*), representing a substantial fraction of the genome. Each odorant receptor recognizes an overlapping set of many structurally distinct odor molecules (*11, 12*). Unlike the case of bitter taste in which all bitter receptors are co-expressed in a bitter cell, vertebrate olfactory neurons express only a single allele of a single odorant receptor gene (*35, 36*). The dimension represented by each receptor is then faithfully transmitted one synapse forward, because axons of all olfactory neurons expressing the same receptor project to a single glomerulus in the olfactory bulb (*15*). Glomerulus input is transmitted to mitral cell output neurons, and relayed to a number of olfactory cortical areas. At these higher levels, there is no evidence for data compression. In fact the current evidence argues the opposite. Most pertinent for olfactory discrimination is the piriform cortex, which selects small groups of mitral cell inputs in a combinatorial way (*16*). Piriform encoding is not only random with regard to input, but also dynamic and highly flexible (*37*), raising the possibility of a vast information coding capacity in olfactory cortex.

Therefore, olfactory sensory neurons expressing the same odorant receptor each represent a dimension corresponding to the receptor’s ligand specificity. Following this logic, the olfactory system is about 400-dimensional in humans, 1000-dimensional in rodents. We shall continue to use the numbers for humans, but note that most of the features have been demonstrated in rodent studies. Thus the extant evidence implies that the dimensionality of olfaction remains as close to 400 as is biologically feasible.

## Dimensional reduction of olfactory data

Faced with this daunting complexity, many researchers in the field of olfaction have used dimensionality reduction procedures to reduce the problem of understanding smell to fewer dimensions, usually from 2 to 10. These approaches are designed to compress complex data into the smallest number of components; they are designed to reduce dimensions and must always do so.

Dimensionality reduction can be divided into two categories.

Those that gather attributes of each odorant independently (*31, 38-45*):

- a set of M monomolecular odorants is considered (occasionally natural mixtures, but never mixtures of the set of odorants themselves)
- a set of S words (sometimes semantic labels, sometimes physicochemical attributes) pertaining to the odorants,
- an MxS matrix of assignments of labels to odorants (either Boolean or graded) is created, either by psychophysical quizzing or by analytical software, and then
- analyzed by a dimensionality reduction algorithm such as Principal Component Analysis (PCA)

Those that compare odorants to each other (*38-40, 46-50*):

- a set of M monomolecular odorants is considered (occasionally complex mixtures, but never mixtures of the set of odorants themselves)
- M(M-1)/2 psychophysical comparisons of perceptual similarity of odorants are made
- a method for converting similarities/dissimilarities to distances is used to create a MxM matrix of distances which is then
- analyzed by an embedding/dimensional reduction algorithm such as MultiDimensional Scaling (MDS)

The need for multiple semantic labels per odorant is telling, because in every other sensory modality there are natural words that describe the perceptual dimensions associated with every cell type, such as “red,” “high-pitched,” “bitter,” or “itch.” Such a procedure is logically unable to discover the dimensionality of olfactory representations in the sense we have used it up to now. Such a method is likely to be uncovering the topological organization of the receptors, the dimensionality of the organization of the system. In other words, it is equivalent to discovering the two-dimensional nature of the retina.

Dimensional reduction methods are neither useful for detecting the true dimensionality of a system, nor, as the word “reduction” in their name implies, are they good at preserving the true dimensionality of smell. This is trivially verified for PCA: analyzing an MxM matrix of independent-identically-distributed-Gaussian numbers, which naively represents an isotropic spherically-symmetric N-dimensional space where all of the N dimensions are equally important, will show 50% of the variability to be concentrated on the first 18% of the eigenvalues and, conversely, the lower 50% of the eigenvalues only account for 10% of the variability. This is because, unless there is an enormously larger number of examples than dimensions, there shall be only one point per dimension to establish the size of the data cloud, and the largest probability of a Gaussian is at zero. Such a distribution of eigenvalues would lead many to believe the dimensionality of the system is far below its true value. Because of theorems showing that the eigenspectrum of Boolean random matrices follows the same laws as those for Gaussian matrices, this result holds as a null hypothesis for matrices associating labels to odorants. Even if the above hurdles were overcome, the dimensionality of olfactory semantic categorization of monomolecular odorants is not germane to the distinct psychophysical task of discrimination of complex mixtures.

An interesting twist into this discussion is brought by the work of (*45*). The authors did not apply a global dimensionality-reduction scheme, but tried instead to map out a lower-dimensional curved manifold within the full space. They found that a curved 2D surface within their larger dimensional space contained all the monomolecular odorants. Intriguingly, they describe their surface as being curled up in all relevant dimensions.

The curvature of the surface means that the line joining any two monomolecular odorants, which describes their combination in a mixture of varying proportions, will be mapped outside of the “monomolecular” surface. The farther away, the more curved the surface or the more separated the elements of the mixture. This organization implies directly that combinations of odorants require more dimensions for their description than the monomolecular odorants on the low-dimensional surface.

## The role of dimensionality: D=25 suffices

The central argument in (*4*) is that our estimate counted stimuli rather than percepts. The implicit assumption that permits us to do so is that stimuli which are sufficiently far from one another evoke percepts which are also far enough from one another. The space of stimuli, being combinatorial, grows very rapidly. If the space of percepts grew more slowly, it would eventually be unable to keep up and the assumption would be broken: multiple far-away stimuli would map onto the same repeated percept.

What do we mean by “grows rapidly” ? In any space in which there is a proper notion of distance, we can define a *sphere of radius R* around a point called its *center*, as the set of all elements of the space which lie at a distance of exactly R from the center; the *ball of radius R* is the set of all points which lie at a distance <= R. In 3D space, the sphere is just a hollow shell of a surface, while the ball is a solid object: the sphere plus all its contents, center included. The volume of the ball of radius R counts the number of elements contained within it.

Imagine a binary tree, and consider traversal of any link as a distance of 1. The sphere of radius R centered at the root of the tree is the level R in the tree, which contains 2^R elements, while the ball of radius R contains all levels up to and including level R, and has a volume of 2^(R+1)-1 elements. Since the balls of radius R in a Euclidean space of dimension D grow like R^D, for any fixed dimension D there will be a sufficiently-large radius R at which the tree grows faster than the Euclidean space. Eventually, there will be no way to accommodate so many elements within the space. Thus spaces that grow rapidly, like binary trees, cannot be embedded in any Euclidean space because no matter how large the dimension, Euclidean spaces *eventually* run out of space. More precisely, the number of stimuli at a distance smaller than R around a reference stimulus S0 grows combinatorially fast, while the number of distinct percepts evoked by those stimuli grows at first slowly and then saturates. In the case of the YumYuk and Circle models in (*4*) this is true trivially, because they have a small discrete number of percepts and they get repeated at large distances in random ways. The simplest nontrivial way in which the exponentially-growing space of combinations can fail to give exponentially many percepts is if the latter is a low-dimensional Euclidean space. The RGB mode presented in (*4*) is such a model and is based on color perception.

The Euclidean character of the space is essential, as there exist numerous examples of *finite-dimensional hyperbolic spaces*, in which spheres grow as exponentially-fast functions of the radius. The most famous of these is the Poincaré disk (or pseudosphere), the simply-connected two-dimensional manifold of constant negative curvature, first introduced by Beltrami in 1868, and familiar to many from numerous illustrations by M. C. Escher. In the Poincaré disk, the circles of radius p enclose an area A(p)= 2n(cosh p - 1), which grows exponentially, even though it is everywhere locally a 2-dimensional space. It is thus important to keep in mind that a perceptual space may be locally of small dimensionality, yet can be globally hyperbolic or even branched, in which case exponentially many percepts may still be accommodated. All open combinatorial spaces have such structure: the space of words in a finite alphabet and spaces of combinations of image elements or sounds.

Our discrimination study did not have to go to arbitrarily long distances, because our spaces have finite radii. In particular the set of combinations of 30 elements, from which the bound of 1 trillion was derived, has a radius of *precisely* 30. There is nothing beyond 30 because you cannot change more elements than you started with. Therefore, even though an infinitely-growing space of combinations will eventually outrun every Euclidean space of percepts, in our case we only need the Euclidean space to keep up with that growth until R=30 (Figure 1). The volume of the sphere of radius 30 grows so fast with the dimension D, basically an extra factor of 10 per each extra dimension such that for dimensions not much larger than 20, Euclidean spaces have exponentially more space than needed. Thus the assertion in (*4*) that the conclusions in our paper required that the dimensionality of olfactory representations to be “at least 128” is incorrect. The 128-dimensional Euclidean sphere of radius 30 is larger than the space of all combinations of (30 out of 128) by a factor of 4x10^106^. Of course, we still need those 25 dimensions to contain the space generated by our combinations, but that is where the choice of 128 elements matters.

**Figure 1.**
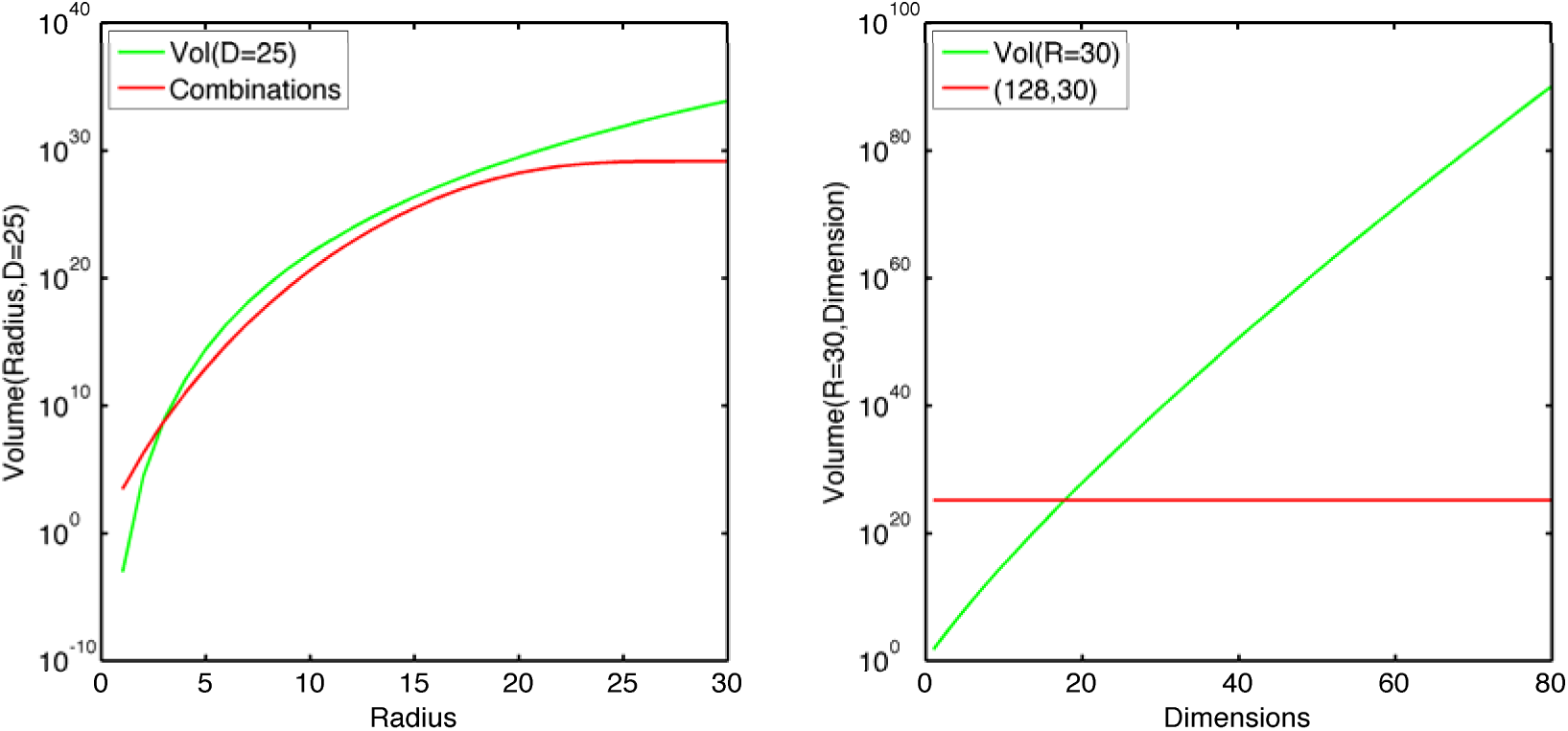
Dimensionality of the smallest Euclidean space that can contain all of the combinations without folding. Left, the volume of a sphere of radius R; red: number of combinations of 30 elements out of 128 having up to R different elements from a reference combination; green, 25-dimensional Euclidean space. For R>3 the 25D Euclidean space can accommodate all the possible combinations without repeating or folding over. At R=30 the number of combinations equals the total number of possible mixtures of 30. Right, the volume of a ball of radius R=30; green, Euclidean space of dimension D; red, the total number of combinations of 30 elements (from the left panel). For even moderately larger dimensions, such Euclidean spaces have astronomically more space than needed to embed our combinations. The code used to generate all figures in this article is available by request from M.O.M.

Having established that the bound to beat, 25 dimensions, is well within reason for a sensory system with 400 independent sensors whose signals are keep strictly segregated, we proceed to study carefully the models presented in (*4*).

## The RGB model

The RGB model in (*4*) supplants a large dimensionality of possible percepts with high discrimination acuity in its 3-dimensional space to generate a million percepts. Let us re-state the model as “generate N vectors of length 1/C, choose C of them and add them up” to “generate N vectors of length 1, choose C of them and average them”, which allows us to choose the same N vectors for tests involving multiple values of C. When one averages C vectors, the average vector is closer to the mean of the distribution from which they were drawn by a factor of sqrt(C). Therefore, the space of combinations of C vectors contains only C^(-3/2) of the original percepts. When a single vector is changed, this average is shifted to another position, and this shift follows the distribution of differences of the vectors. In any one coordinate, the probability of a shift of magnitude S diminishes linearly from a maximum at S=0 to a minimum at S=1/C, making small shifts more likely, but in three coordinates the probability distribution of the max distance is zero at 0 (with a 3/2 power law) and zero at 1/C with an average of 0.5432/C. When 0.5432/C is about the limen, then a single shift of 1 vector is likely to make the changed mixture discriminable, and since 0.54/30 > 0.01 this means mixture changes of 1 component in 30 are discriminable within this model. On the other hand, the effect compresses at large numbers of changed components by a square-root effect. Changing D vectors only moves you about sqrt(D)/C in the space, until the moment you change all C vectors and you are back to the original distribution. This forces the discriminability curve to have the outline of a square-root-like function, rather steep at the origin where the maximal differential discrimination is achieved by change of a single vector, and leveling off rapidly.

Please note that the overall curve depends on the initial choice of 128 vectors, as shown in Figure 1, because the amount by which vectors do move when components change depends on the *empirical* standard deviation of the vectors, not on the asymptotic, and this empirical standard deviation fluctuates by over 10%. As a result of the rapid leveling off of the discriminability curves, they fail to approach 100% at large changes. For instance, for C=30 the last points (D=30) approach 97%, meaning 3% of the stimulus pairs of 30 components without any component in common are not discriminable Figure 2.

**Figure 2.**
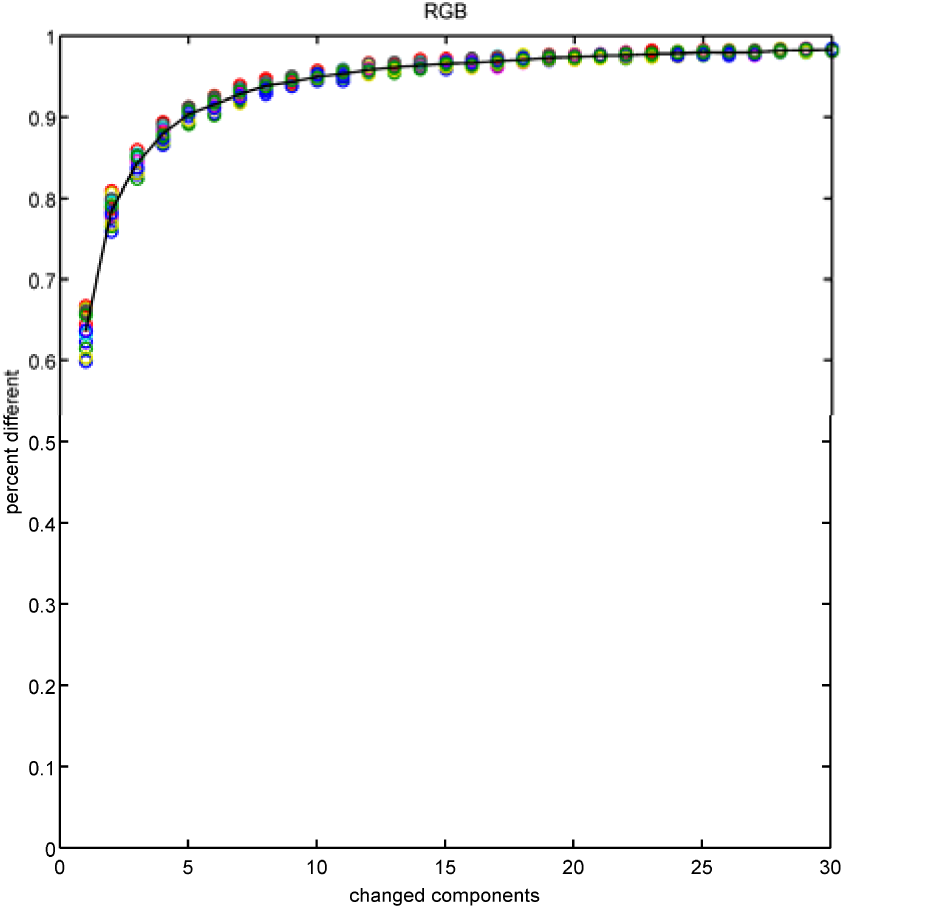
Discriminability curve of the RGB model for C=30, generated from simulated psychophysical comparisons using the method described in (*4*). Each circle is the result of comparing 10,000 different pairs of stimuli by 11 subjects. The 16 different-colored circles represent 16 choices of the 128 basis elements, showing how much the entire curve shifts up or down depending on this random choice. The black curve is the average value over the 16 choices of components. Please note that the rightmost point is 3% away from being 100%, keeping in mind that there is nothing to the right of this curve – 30 components changed corresponds to zero overlap. This is because the set of combinations of 30 components does not cover the million percepts of the entire space, but a much smaller set. NOTE: The model as outlined in (*4*) is insufficiently specified for direct reproduction of its Figure 2B. For instance, all basis elements are made of the same “length” without specifying what the length definition is (L1 norm, L2 norm, sRGB luminance?). The comparison between the noise-corrupted stimuli to simulate the psychophysics is also carried out using an unspecified distance. We have chosen for these parameters what appear to be reasonable guesses – L1 norm set to 1 for length definition, L2 norm for psychophysics comparisons, and we computed the amplitude of the Gaussian noise that generates a limen of 0.01 in either direction, unstated in (*4*), to be approx. 0.5698*0.01.

To extend the RGB model to D dimensions we will ask for what values of D is the perceptual space large enough to hold all of the combinations we’re discussing? Of course, given a small enough discrimination limen this will happen for any D, so we need to adjust the limen to satisfy some form of benchmark. So, for every dimension D, we shall search for the value of the limen such that the threshold for 50% recognition in mixtures of 30 components is 15 components changed, the threshold that gave rise to our estimate of one trillion (*1*).

As the dimensions increase this limen L slowly increases, as shown in Figure 3 (left). A computation of the total number of mutually-discriminable percepts available in the model would say it is roughly L^{-D}, the total number of little hypercubes of edge L in the unit hypercube. As Figure 3 (right) shows, the curve for this total number of percepts rises exponentially with dimension D, at a rate of roughly 8-fold increase per unit dimension in the range we are considering, and crosses the one trillion value right next to D=12.

**Figure 3.**
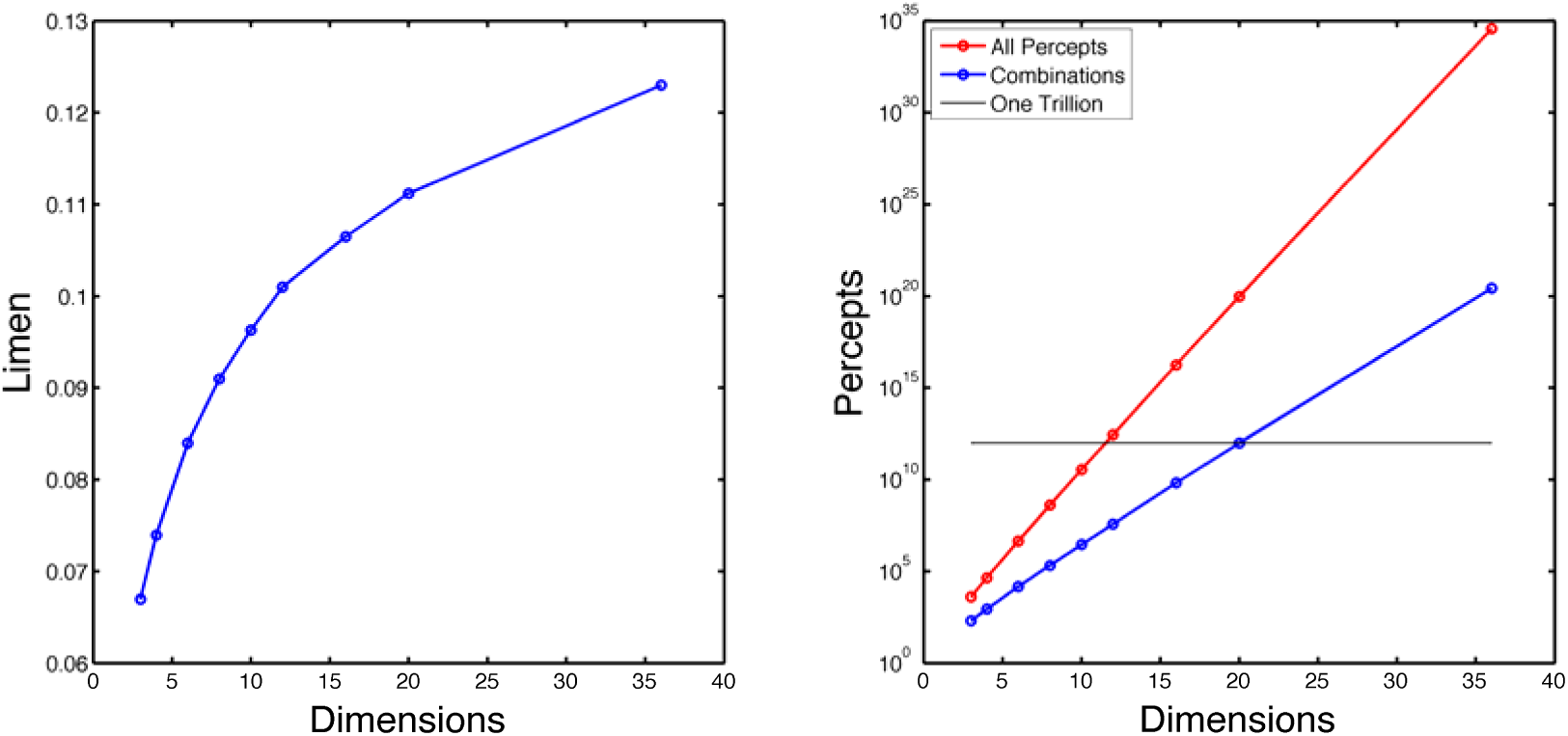
Left, limen of discrimination as a function of dimension, for the D-dimensional RGB model to have a discrimination threshold at 15 components changed for 30-component mixtures. Right, the total number of percepts in the model (blue) and the number of percepts in the region mapped by combinations of 30 components (red).

However, this would be the right answer for the wrong reasons, because 95% of combinations of C components, in this model, map within a smaller hypercube centered within the larger one, with an edge of roughly 1.8/sqrt(C); C=30 in our case. We would like to also know how many distinguishable percepts there are in this exponentially smaller region. The answer, the blue curve in Figure 3(right), crosses the trillion threshold at D=20. At this dimensionality, the RGB model stops repeating percepts for the different combinations of 30 vectors that are further away than 15 changes.

To simplify our calculations, we will avoid the normalization step, so the basis of 128 vectors will be unnormalized, and instead of simulating the psychophysics we shall simply declare two stimuli to be distinguishable if any of their coordinates differ by more than a limen (so distance is measured in the Linfty metric).

## The Circle model

In the Circle model proposed in (*4*), N vectors on the unit circle are chosen, and combinations are evaluated by averaging the components *as complex numbers* and re-normalizing them by projecting the average back to the unit circle. When C unit vectors are averaged, the average most likely falls within a disk of radius sqrt(C) around the origin. In the Circle model this distribution is then projected back onto the unit circle. When the average of the C vectors is sufficiently close to the origin, any small change can shift the projection into a completely different quadrant of the circle.

This simple conceit permits the Circle model to generate rapid and random changes of percept for small numbers of changes in components. However, this also exacerbates the dependence of the discriminability results on the choice of 128 vectors. In particular, if the *empirical average* of the 128 vectors is sufficiently far away from the origin, the probability distribution of the percepts associated to combinations of 30 components is far from flat, generating *de facto* fewer than the expected number of percepts, which for the stated noise amplitude of 0.4 is 10.7. As a result, our simulations show (Figure 4) that for 30 changed components the discriminability curves never reach the naïve expectation of 90%, with one out of 16 simulations reaching 72%. This falls far short of our data.

**Figure 4.**
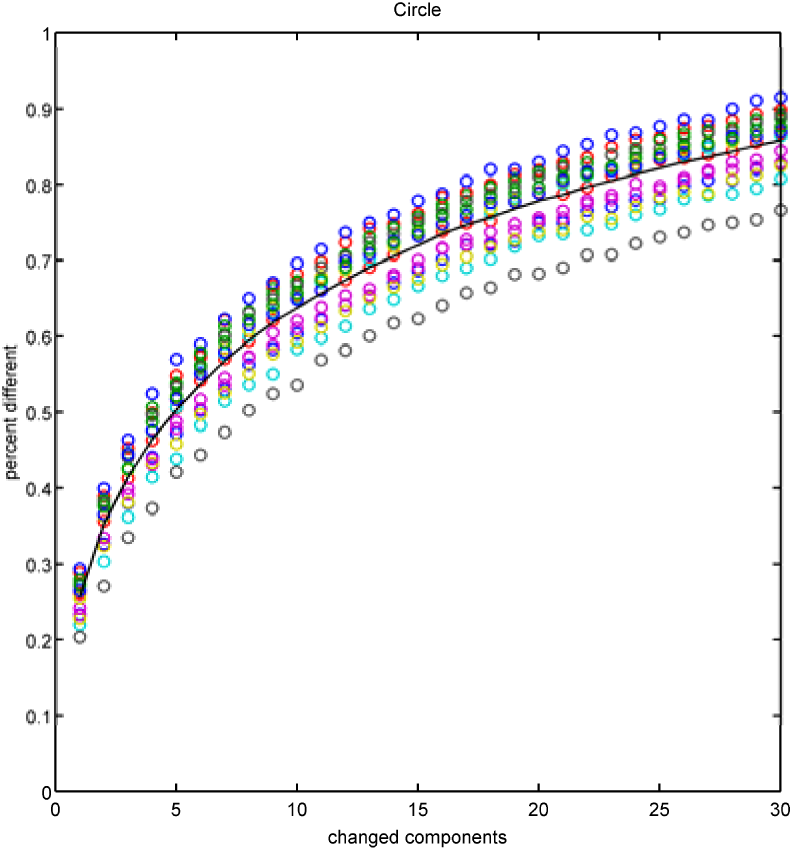
Discriminability fraction of the Circle model, as defined in (*4*). Notice the convex outline and the large fluctuations for different random choices of the 128 elements. Depending on how far away from the origin is the average of the 128 basis elements, the number of *effective* percepts generated by random choice of 30 elements is fewer than the theoretical average of 10.7 percepts for the value of the noise amplitude used (0.4), leading to a *mean* value of 84.7% at no overlap.

## The YumYuk model

The analysis in (*4*) claims that applying our procedure to the YumYuk model would give an estimate of millions of percepts when there are in fact only three. But there are problems with this model. Our experimental setup created mixtures from a set of 128 mutually-discriminable component odors (*2*). Within YumYuk, any single component only elicits *Meh*. In fact, any combination of two components only elicits *Meh*, as the absolute value of the sum of valences has to exceed 2 to elicit either a *Yum* or a *Yuk*. This threshold of 2 has been chosen clearly so that the discrimination curves for 30 components cross 50% at a fair number, but interestingly the model simply does not work for other components. The discrimination curve for 10 components never crosses 50%, and the discrimination curve for 60 components regresses back, so the threshold is non-monotonic (Figure 5). Therefore, the statement that our analysis method applied to YumYuk would estimate millions of percepts, rather than 3, is grounded in a flawed model.

**Figure 5.**
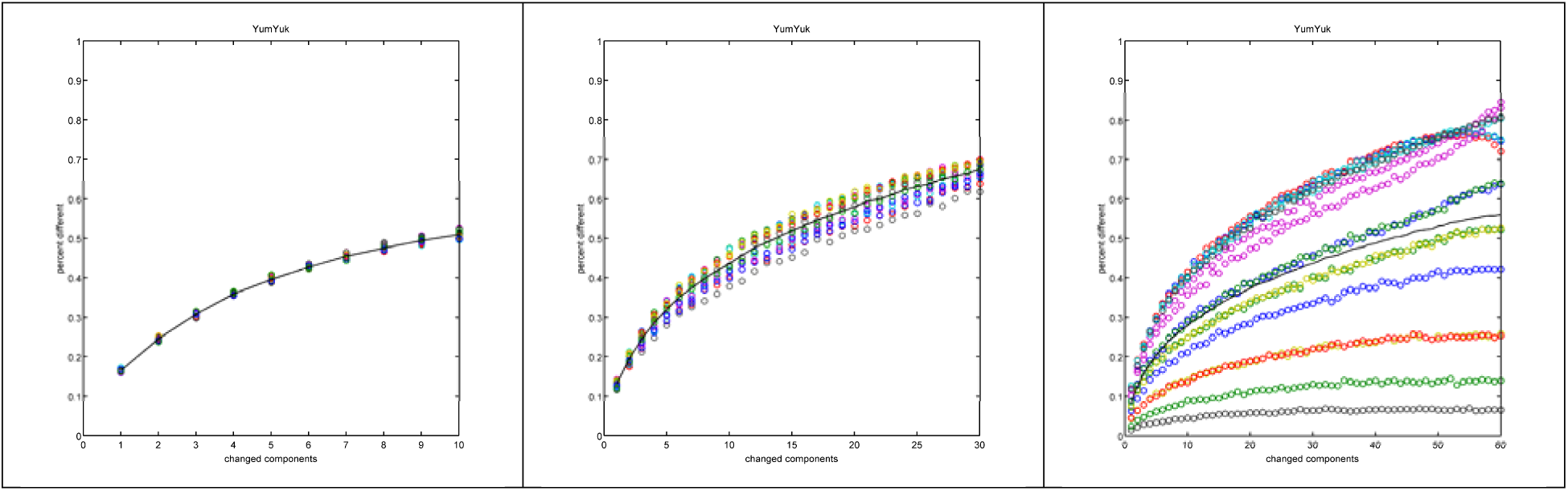
Discriminability curves for the YumYuk model. Circles are the outcome of 10,000 comparisons of two stimuli, and each experiment was repeated 16 times for different choices of the 128 single components. Left, for C=10 the curve never crosses the threshold, so *we would have estimated that the number of mutually discriminable combinations of 10 is 2*, which is in fact accurate as the Meh percept is more probable than Yum or Yuk for C=10. Middle, C=30, the curves do cross threshold at 14 components as stated in (*4*). However, the rightmost point, at 30 changed components, only shows 67% discriminability, leading directly to an estimate that there are no more than 3 percepts in this space. Right, as the number of components becomes larger (C=60,), the curves are dominated by the empirical average of the basis, with some experiments never crossing threshold and others doing so early, as the Meh percept becomes less probable that Yum or Yuk.

## Conclusions

*The problem of odor classification has been historically approached by an almost fruitless series of attempts at categorization. Relating our psychological experience to the nature of the stimulus has been made doubly difficult because, in addition to not knowing what the stimulating process is, there has been little agreement about the description of the odor experience.*

*If a psychophysics is to be established in olfaction, it would seem imperative to illuminate more clearly the nature of our olfactory experience.*

M. H. Woskow, *Multidimensional Scaling of Odors* (1966) (*46*).

In this paper we have carefully considered the mathematical arguments raised in (*4*), and find that they do not stand up to the biological evidence that olfaction is a high-dimensional sense. A simple geometric calculation shows that to avoid the mathematical problems pointed out in (*4*), the dimensionality does not need to be 400, the number of functional human olfactory receptors (*7, 8*), or even 128 as claimed in (*4*), for in fact D≥25 suffices. Calculations based on the RGB model in (*4*) show that for our assumptions to hold we need D≥12 to D≥20 depending on details. Since all experimental evidence points to olfaction being a high-dimensional sense, and nothing in (*4*) has made us revise this belief, our confidence in the correctness of our results is entirely unchanged and we stand by our original estimate.

We have shown at length why the argument in (*4*) is unsound. First, the evidence for dimensional segregation in the 400 D range does not only include receptor diversity, but also expression of single receptors in each sensory cell, axonal segregation for targeting onto the olfactory bulb, and dimensionality expansion as piriform cortex target neurons sample mitral cell input. Second, the studies invoked to counter the claims of high dimensionality in olfaction have no scientific connection to our task. These studies estimated the rank of a matrix associating monomolecular odorants to semantic labels, thereby studying the association between single stimuli and categorization. Our work ignores semantic categories of smell and instead probes olfactory discrimination.

The debate between our work on olfactory discrimination (*1*) and the critique (*4*) raises a few additional interesting questions. First, what is olfaction “for” ? Does it exist to place percepts into semantic categories represented by everyday words such as “rose,” “vanilla,” or “chocolate” ? Or to arrange percepts on a valence axis from “extremely unpleasant” to “extremely pleasant” as the YumYuk model proposes? Or to make fine discrimination judgments between perceptually similar odors? In practice, the human sense of smell is capable of all of these tasks, and all are equally important for our conscious experience of odorants. Second, does the olfactory system compress information or expand it? The proposal in (*4*) is that the extraordinarily large number of odorant receptors exists merely to capture information about a large number of olfactory stimuli, and then compress them into a small number of semantic categories. Although not definitively proven, neither the circuit organization of the olfactory system, nor existing psychophysical evidence supports this view.

One suggestion in (*4*) is that the way forward is to study this problem in mice. Indeed, there have been great advances in genetically-guided optogenetic activation of olfactory circuitry in mice (*37, 51*). Measuring olfactory discrimination in mice requires hundreds of trials of operant conditioning per odor pair (*52*). Contrast this to naïve human subjects, who can carry out >50 discrimination trials per hour with no prior training (*1, 22*).

Therefore the number of odor pairs that can realistically be tested in mice is orders of magnitude smaller than in humans. Even using discrimination testing in humans, it is obvious that an extrapolation of some kind must be employed. The number of all possible odorants is unknown, and it is not feasible to carry out trillions of pairwise discrimination tests.

The field of olfaction will need to overcome a number of important challenges if further progress is to be made. We need clarity on the transformation of the chemical structure of an odor to its percept; the ligand selectivity of all human odorant receptors needs to be defined; how olfactory cortex process the information relayed from the olfactory bulb must be understood more clearly; and how olfactory stimulus space relates to olfactory perceptual space in the unknown dimensions of smell must be solved.

The question of the dimensionality of smell is an important one, and we hope that the discussions started by our paper stimulate more efforts to make progress in this area.

## Acknowledgments

We thank Cori Bargmann, Stuart Firestein, and Avery Gilbert for helpful comments on the manuscript; and Larry Abbott, Richard Axel, Bill Bialek, Linda Buck, Jeff Friedman, Markus Meister, Jonathan Victor, and Torsten Wiesel for lively discussions.

## Conflict of interest

L.B.V. is a member of the scientific advisory board of International Flavors and Fragrances. M.O.M. and A.K. declare no conflicts of interest.

